# Automated Surface-Based Segmentation of Deep Gray Matter Regions Based on Diffusion Tensor Images Reveals Unique Age Trajectories Over the Healthy Lifespan

**DOI:** 10.1101/2023.10.04.560912

**Authors:** Graham Little, J. Alejandro Acosta-Franco, Christian Beaulieu

## Abstract

Many studies have demonstrated unique trajectories of deep gray matter (GM) volumes over development and aging, suggesting but not measuring microstructural alterations over the lifespan. Only a few studies have measured diffusion tensor imaging (DTI) parameters in deep GM or reported these values across a wide age range in a large cohort. To enable efficient DTI studies of deep GM in large cohorts without the need of T1-weighted images, an automated segmentation technique is proposed here that works solely on parametric maps calculated from DTI. The algorithm segments the globus pallidus, striatum, thalamus, hippocampus and amygdala per hemisphere by deforming 3D models of these structures to their boundaries visible on the contrast provided by diffusion tensor maps and images alone. This new DTI-only method is compared against standard T1-weighted image segmentation for (i) 1.25 mm isotropic diffusion data from the Human Connectome Project (HCP) test-retest cohort (n=44) and (ii) 1.5 mm isotropic test-retest diffusion data from a local normative study (n=24). Dice coefficients of voxel overlap between methods in the HCP test-retest cohort were high (>0.7) for 7 of 10 structures, but were low for the left globus pallidus (0.54) and left/right amygdala (0.67, 0.69). The proposed DTI-only segmentation qualitatively appeared more accurate and yielded smaller volumes than T1w for 8/10 structures in both cohorts, with the exception of the globus pallidus which showed larger volumes in the HCP test-retest data but lower volumes in the local normative study data. The DTI-only segmentation method was then applied to two local single site development/aging ‘lifespan’ cohorts (cohort 1: n=365 5-90 years, cohort 2: n= 164 5-74 years) to assess age changes in volume, fractional anisotropy (FA) and mean diffusivity (MD). In both cohorts, MD trajectories were quadratic for all five structures, decreasing slightly and then increasing after ∼30-35 years. In cohort #1, FA trajectories remained flat from 5 to ∼25 years and then started to decrease for the globus pallidus and hippocampus and over 5 to 90 years, FA decreased linearly for amygdala, increased linearly for striatum, and remained constant for the thalamus. In the second cohort, using an alternate acquisition protocol, the FA trajectories of all 5 structures across all ages were similar, except for the globus pallidus and thalamus which both increased in value from 5 ∼ 20 years and likely reflect differences in acquisition details. Notably, the development and aging trajectories for DTI were distinct from those of the deep GM volumes. The proposed automated deep GM segmentation method on DTI-only will facilitate the analysis of deep GM DTI (currently ignored in nearly all studies despite the data there within the field-of-view) and will be advantageous particularly for studies that do not have a T1-weighted scan, as in many clinical populations.

## 1. Introduction

Magnetic resonance imaging (MRI) studies of subcortical/deep gray matter (GM) structures (e.g. globus pallidus, caudate, nucleus accumbens, putamen, thalamus, hippocampus, amygdala) have primarily focused on their volume (measured on T1 scans) that show unique non-linear cross-sectional trajectories over the healthy lifespan that increase until adolescence, peaking between ∼15-25 years of age and thereafter decrease with aging (e.g. Bethlehem et al., 2022; Fjell et al., 2013; Narvacan et al., 2017). There are far fewer investigations of deep GM using diffusion tensor imaging (DTI) which have provided complementary parameters indicative of region-dependent microstructural changes of multiple deep GM regions with typical child development to young adulthood (e.g. Lebel et al., 2008; Mah et al., 2017; Snook et al., 2005) and aging (e.g. Bhagat & Beaulieu, 2004; Pal et al., 2011; Pfefferbaum et al., 2010; Wang et al., 2010). In contrast to DTI of white matter, a primary reason for so few DTI studies of deep GM in large healthy populations is the limitations of commonly used analysis methodologies, such as manual region-of-interest (ROI) segmentation that is time consuming. For practical limitations, such manual ROI studies then usually extract measurements from only a portion of the structures such as on a single slice.

There are two main automated approaches to segment the deep GM to extract diffusion parameters. The first approach uses a set of pre-segmented ROIs in a standard diffusion template space (often age-relevant) that is then applied to parcellate brain regions in coregistered diffusion images and maps within individual participants (Oishi et al., 2013; Zhao et al., 2021). However, this relies on proper registration which can be problematic, particularly for small deep GM regions, given differences in the anatomical shape and image acquisition (e.g. in-plane resolution, slice thickness, image distortions, etc.). Furthermore, the DTI templates (e.g. ICBM-DTI-81) are usually focused on parcellations of white matter regions (Mori et al., 2008). The second approach involves identifying voxels belonging to deep GM regions using one of many automated methods on an additionally acquired T1-weighted (T1w) structural image that is then co-registered to the diffusion images (Mah et al., 2017). However, this registration is prone to errors because the diffusion images acquired with single-shot EPI typically have spatial distortions not present in the structural image and are usually acquired at a much lower spatial resolution. In addition, basing segmentation techniques on T1 weighting alone is problematic because many regions adjacent to deep GM have poor WM/GM contrast; for example, the lateral boundaries of the thalamus are most often indiscernible on T1 weighted images. This lack of contrast limits the use of deformable mesh based segmentation methods that use the 3D geometry of the target structure to align the edges of a 3D model to regions with large intensity gradients on an image, as has been developed for cortex (Dale et al., 1999; Kim et al., 2005; Schuh et al., 2017). To employ deformation-based segmentation methods, FA maps, which have excellent WM/GM contrast in regions surrounding the basal ganglia, have been proposed to improve mesh-based segmentation compared to methods using a single image contrast (Visser et al., 2016). Given that the other boundaries of the subcortical GM are also visible on diffusion weighted images and maps, this suggests that diffusion MRI alone may provide the contrast necessary to segment the deep GM structures as has been achieved recently for the cortex (Little & Beaulieu, 2021), avoiding the need for additional structural imaging, or problematic registration between different image types.

To date, only two studies have developed deep GM segmentation methods that work on native diffusion images with both techniques relying on deep learning. The first approach used diffusion parameter maps (axial diffusivity, mean diffusivity, radial diffusivity, and apparent fiber density) as input to learn a 10 structure segmentation directly on native diffusion images yielding moderate Dice-scores between 0.63 and 0.80 compared to FreeSurfer T1-weighted image segmentations applied to registered diffusion images (Theaud et al., 2022). However, the focus was not on extracting diffusion parameters from deep GM, but rather to improve anatomically constrained tractography of WM tracts. The second approach generated synthetic T1-weighted images from diffusion data and then applied a standard T1w segmentation approach (e.g. FreeSurfer) to delineate various brain structures (Li et al., 2023). When compared to segmentations generated on the original T1w images, the segmentations based on the non-synthetic images showed significant overlap indicated by Dice-scores ranging from 0.73 to 0.92 for separate deep GM structures. However, synthetically reconstructed T1w images also contain the same poor WM/GM contrast adjacent to deep GM structures that is apparent on native T1w images, hence this method does not utilize a clear advantage DTI has over other imaging modalities for identifying the deep GM boundaries adjacent to the internal capsule.

The purpose here is to develop an automated segmentation approach for deep GM that works solely on diffusion tensor images and maps that can be calculated from a large variety of typically acquired diffusion protocols. This technique avoids potential for erroneous registration with other scans like T1w, as well as not needing a T1w image in the first place as many clinical protocols do not acquire it. This is a natural extension of a previously developed surface-based segmentation method for the cerebral cortex that also only relies on the contrast available from diffusion tensor images and maps (Little & Beaulieu, 2021). The new method is first validated based on segmentation overlap using Human Connectome Project test/retest 1.25 mm isotropic diffusion data and 0.7 mm isotropic T1w images, with both modalities registered and corrected for distortions.

Second, the method is evaluated on locally run 1.5 mm isotropic whole brain diffusion acquisition acquired in ∼3 minutes in a test/retest cohort comparing regional brain volumes between the new method and FreeSurfer segmentations output from T1w images. As a proof of concept while also yielding the first DTI healthy “lifespan” study across multiple deep GM structures, this novel method was used on 1.5 mm isotropic diffusion MRI data to evaluate age changes of diffusion in the striatum, thalamus, globus pallidus, hippocampus, and amygdala in two cross-sectional cohorts of typical individuals (N=357 and N=164) across much of the lifespan (5-90 years).

## 2. Materials and Methods

### 2.1. Participants and Data Acquisition

The method described in Sections 2.2/2.3 below was applied to high-resolution 1.25 mm isotropic diffusion data (already coregistered to the structural images) from a test-retest cohort (n=44; age 30 ± 3 (22 - 35) years, 31 females, scanned 135 ± 63 (18 - 328) days between test-retest sessions) from the Human Connectome Project (HCP) since it already includes subcortical segmentations generated with a standard FreeSurfer pipeline using high resolution 0.7 mm isotropic T1 and T2 weighted images. Note that only the b0 (5 averages) and b1000 (90 directions) acquired twice, in L/R and R/L phase encode directions (Sotiropoulos et al., 2013) were used in the current study.

The local healthy participant study was approved by the Human Research Ethics Board at the University of Alberta with written informed consent (and assent for children). First, test-retest diffusion data were acquired from 24 healthy participants (29 ± 8 (20 - 48) years; 13 females) on a 3T Siemens Prisma (64 channel head and neck coil) as part of the University of Alberta normative brain imaging database (Treit et al., 2023). Diffusion tensor imaging was acquired with a single-shot EPI spin-echo sequence: multi-band=2, GRAPPA R=2, 6/8 partial Fourier, 10 b0 s/mm^2^, 6 b500 s/mm^2^ (not used here), 20 b1000 s/mm^2^, 64 b2500 s/mm^2^ (not used here), TR=5160 ms, TE=67 ms, FOV=220 mm, 96 1.5 mm slices with no gap, 1.5×1.5 mm^2^, reconstructed over the 64 channels using sum of squares, with an acquisition time of 9:29 min (or 2:51 min for only the b0 and b1000 shells used for this study). For comparison to more standard T1-based coregistered to DWI segmentation approaches, whole brain T1-weighted imaging was acquired with an MPRAGE sequence: TE/TR/TI=2.4/1800/900 ms, 0.87 x 0.87 x 0.85 mm^3^ resolution, 208 sagittal slices, and acquisition time of 3:39 min.

To assess age related DTI parameter trajectories in 6 deep GM structures, diffusion data were acquired from two separate lifespan cohorts. The first cohort (cohort #1), included 382 subjects recruited under the University of Alberta normative brain imaging database (Treit et al., 2023) and were scanned with the same 1.5 mm isotropic diffusion imaging and T1-weighted MPRAGE protocols as the test-retest cohort above. Amongst these individuals, 365 (38 ± 23 (5-90) years; 209 females) completed adequate diffusion and structural imaging for the proposed analysis. The second cohort (cohort #2), separate from the normative database above, was used to assess the reproducibility of the age related volumetric and DTI trajectories generated in the first cohort. For this cohort 180 subjects were recruited, from which 165 (30 ± 19 (5 - 74) years; 98 females) completed adequate diffusion and structural imaging for the proposed analysis. The lifespan reproducibility cohort used a slightly different diffusion imaging protocol albeit at the same 1.5 mm isotropic acquisition. Diffusion images were acquired on the same Siemens 3T Prisma with a single-shot EPI spin-echo sequence: multi-band=2, GRAPPA R=2, 6/8 partial Fourier, 6 b0, 30 b1000 s/mm^2^, 30 b2000 s/mm^2^ (not used here), TR = 4700 ms, TE = 64 ms, FOV = 220 mm, 90 1.5 mm slices with no gap, 1.5 × 1.5 mm^2^ zero-filled to 0.75 × 0.75 mm^2^ in-plane, reconstructed over the 64 channels using sum of squares, and a 6 min total acquisition time (although only 3.5 min for just b0 and b1000 used for the proposed analysis method). The same T1-weighted MPRAGE protocol was also acquired as above.

### 2.2 DTI Pre-processing

The HCP test-retest diffusion weighted images had already been preprocessed using the HCP recommended pipeline (Glasser et al., 2013). Preprocessing of the University of Alberta diffusion data (test-retest and both lifespan cohorts) was performed with a custom pipeline designed such that only the diffusion images were used as follows. To generate consistent brain masks that were inclusive of all brain tissue voxels, initial tensor models were first fit on unprocessed diffusion data for each subject (Garyfallidis et al., 2014) (DIPY v1.0) and a brain mask was generated on the outputted MD map (Smith, 2002) (BET, FSL v6.0). These tensor maps were recalculated after the following preprocessing steps. Note that for the locally acquired diffusion data, it was important to correct for the non-central chi-squared signal distribution resulting from the sum of squares image reconstruction of the 64 channel phased array data, which was transformed to a gaussian signal distribution using an automated method for estimating the non-central chi-squared distribution model parameters (St-Jean et al., 2020). Diffusion data were then denoised using the non-local spatial angular matching algorithm (St-Jean et al., 2016), corrected for Gibbs ringing (Kellner et al., 2016) and corrected for motion and susceptibility induced / eddy current distortions (Andersson & Sotiropoulos, 2016)(eddy, FSL v6.0).

For all datasets (including the HCP and locally acquired diffusion data) the final tensor models were fit (DIPY v1.0) outputting FA, MD, and primary eigenvector maps. Additionally, a mean b1000 diffusion weighted image (DWI) was output and was corrected for intensity bias using N4 correction based on the spatial variations of signal intensity estimated from the first b0 image (Tustison et al., 2010). A representative mean b1000 DWI is displayed along with FA and MD maps in Figure 1A-C. The mean b1000 DWI emphasizes the boundary of the globus pallidus, whereas the FA and MD maps clearly delineate the remaining boundaries of the striatum, thalamus, and hippocampus/amygdala region. In fact, WM and CSF voxels adjacent to deep GM can be classified accurately based on FA and MD alone (Hasan et al., 2007). A composite FA + MD map was created for each subject by adding the FA map to the MD map (multiplied by 1000) creating a single image that clearly discriminates GM from surrounding WM or CSF regions (Figure 1D).

**Figure 1.**
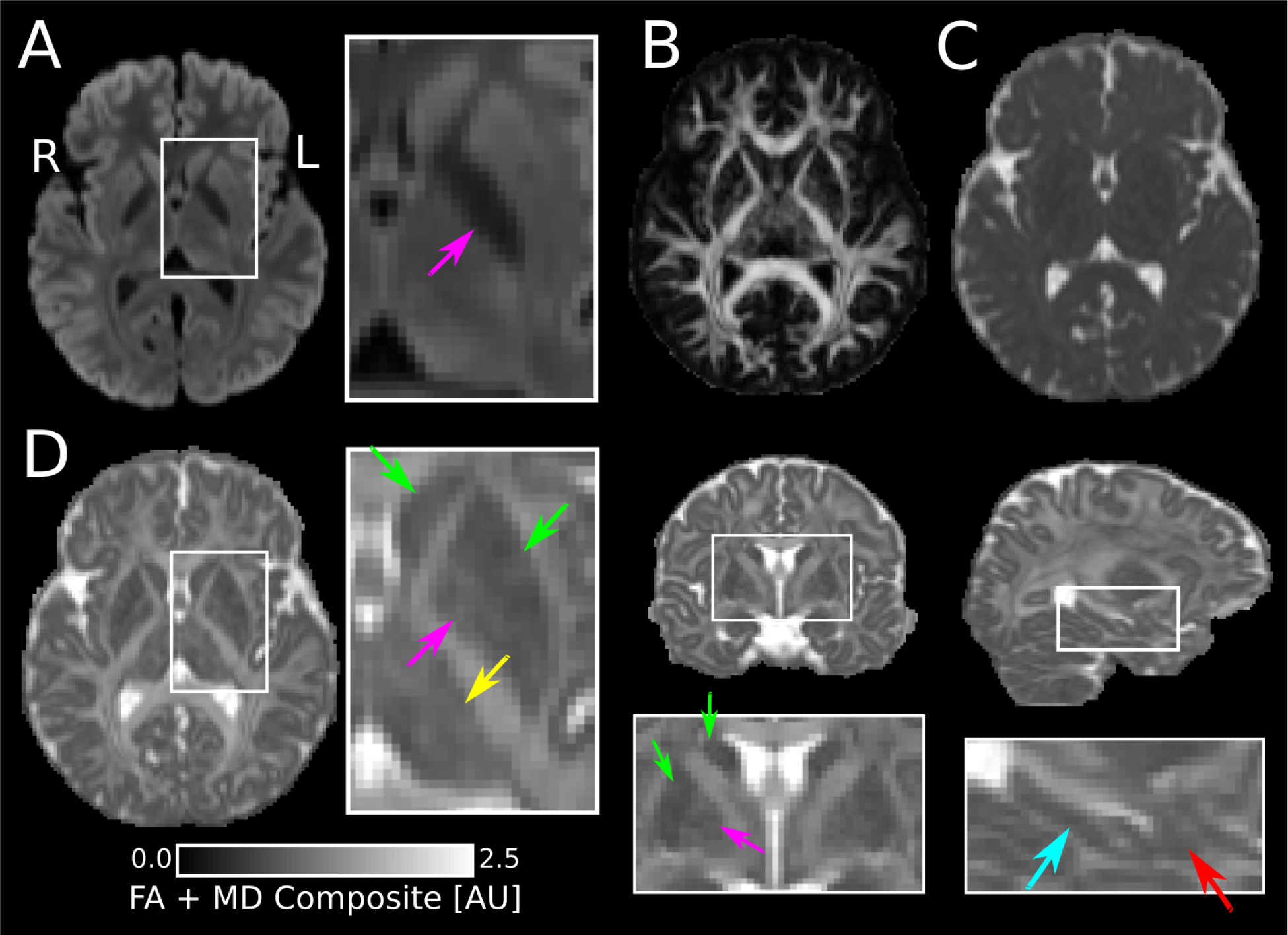
(A) Mean b1000 DWI, (B) FA map and (C) MD axial maps acquired with 1.5×1.5×1.5 mm^3^ resolution at 3T and (D) FA+MD(x1000) composite map in 3 orthogonal orientations. The mean b1000 DWI emphasizes the boundary of the globus pallidus (pink arrow). The FA map clearly delineates the boundary between deep gray matter structures and adjacent white matter, whereas the MD map emphasizes the boundary between the deep gray matter and CSF. When these maps are summed, the FA + MD composite map clearly delineates the boundaries of the striatum (green arrow), thalamus (yellow arrow), hippocampus (cyan arrow) and amygdala (red arrow).

### 2.3 Surface-Based Deep Gray Matter Segmentation on DTI

Typical automated segmentation algorithms that utilize whole brain T1-weighted images aim to segment 7 separate subcortical GM structures in each hemisphere, namely the putamen, caudate, nucleus accumbens, globus pallidus, thalamus, hippocampus and amygdala. However, in our proposed method the contrast on the FA + MD composite map is not sufficient to delineate the substructures of the striatum (nucleus accumbens, putamen and caudate) and to separate the hippocampal head from the amygdala. Thus our proposed method aims to segment 5 subcortical structures namely the striatum, globus pallidus, thalamus, hippocampus and amygdala, where the striatum and hippocampus/amygdala regions are segmented based on the contrast available on the FA + MD composite map. To separate the hippocampus from the amygdala, a secondary segmentation step is performed based on expectation maximization segmentation (ANTs Atropos (Avants et al., 2011)). The entire diffusion only segmentation algorithm is as follows.

The Harvard-Oxford probabilistic atlas was registered to native DTI imaging space by calculating a non-linear transformation between the FSL HCP FA template and the subject’s FA map (ANTS SyN registration (Avants et al., 2008)) per participant extracting left/right probability maps for the caudate, putamen, globus pallidus, nucleus accumbens, thalamus, hippocampus and amygdala. Initial voxel labels for each structure were generated by thresholding by a predetermined value to a singular cluster of voxels for each of the following 4 regions for the left and right hemispheres separately: i) the globus pallidus label was created by thresholding the corresponding probability map by 50%; ii) a striatum label was created by combining the caudate, nucleus accumbens and putamen probability maps and thresholding by 15%; iii) a thalamus label was created using a threshold of 50%; and iv) a hippocampus/amygdala region was created by thresholding hippocampus and amygdala probability maps by 35%. The different probability percentages were selected here to make sure boundaries of the initial structures lay close to the target boundary on most subjects. To remove any voxels containing primarily CSF or WM, voxels were removed from the labels with MD > 1.5 x 10^-3^ mm^2^/s or FA > 0.5, which are tensor parameter values outside of the range for GM (Hasan et al., 2007). Each label was then converted to a surface using the medical image registration toolkit (MIRTK) outputting 3D triangulated meshes consisting of vertices and edges.

Initial 3D meshes were then deformed (MIRTK, deform-mesh) to image edges (regions of large signal change) defined by the mean b1000 image for the globus pallidus, and FA + MD composite map for the remaining regions. Surface deformation algorithms (e.g. cortex segmentation or FSL MIST for subcortical structures) deform vertices towards image edges while constraining surface smoothness. In general, this is accomplished by iteratively moving the vertices based on the external forces (e.g. distance from image edge) and internal forces to regulate surface shape (e.g. curvature at a given vertex or proximity of a vertex to other vertices of the 3D model). For more information about external and internal forces see (Schuh et al., 2017). To accomplish native DTI deep GM segmentation, each structure is deformed separately using a combination of 2 independent external forces based on image edges (from mean DWI or FA + MD composite map) and an implicit force generated for each structure based on the previously registered probabilistic atlas (atlas-based force). For each region the atlas-based force is calculated by adding the probability maps of adjacent GM structures and WM for the probabilistic atlas and subtracting the probability map for the structure being deformed. An example of the atlas based force is shown for the thalamus in Figure 2. This force essentially creates an inward force (positive values) and an outward force (negative values) such that the deformation algorithm will be constrained to image edges on the mean DWI or FA + MD composite map nearby the structure being deformed. Internal forces used here restrict the normal and gaussian curvature of the surfaces and are set to the same value for each structure. Values and details for specific force parameters are provided in the accompanying code available at the first author’s github page (https://github.com/grahamlittlephd/MicroBrain available upon publication).

**Figure 2.**
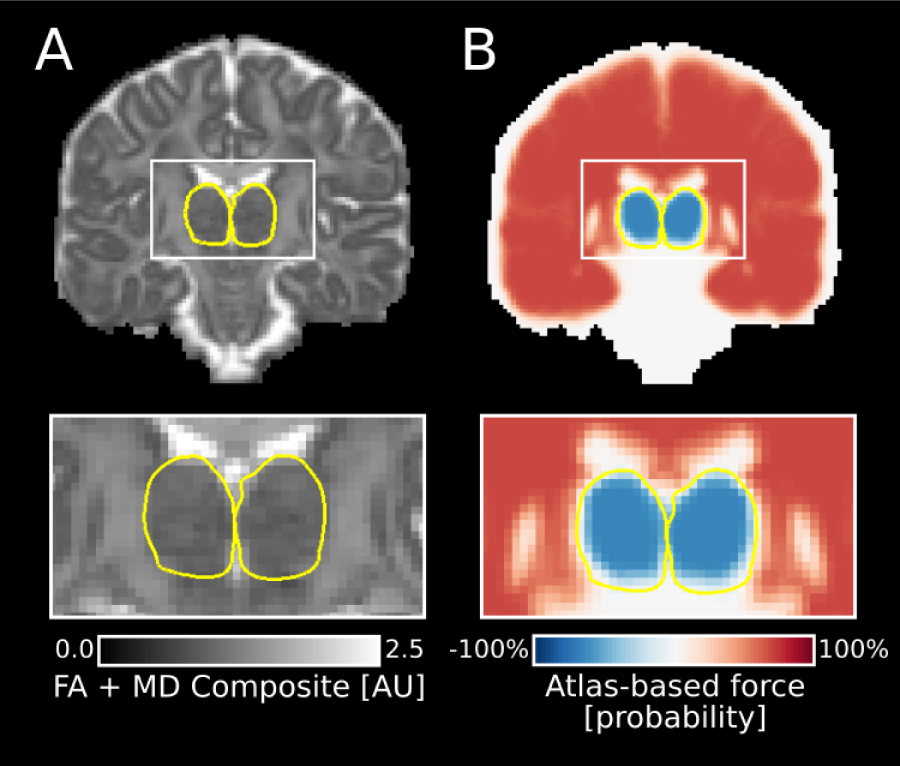
Slice intersection of a 3D thalamus segmentation (yellow) generated using the proposed method shown on the (A) FA + MD composite map and (B) atlas-based force generated from registered probabilistic atlas, where negative values (blue) indicate an outward facing force and positive values (red) indicate an inward facing force. The segmentation algorithm utilizes both forces and in the case of the thalamus will use the FA + MD map where the boundary is clearly defined in the superior and lateral regions of the structure, whereas the atlas-based force is needed to find the inferior edge of the structure.

The segmentation algorithm is visualized in Figure 3 and is summarized as follows. Initial segmentation models are used as input into the deformation segmentation method (Figure 3A) and are then deformed using the atlas based force and the image contrast on either the mean b1000 DWI image or the FA + MD composite map depending on the structure. The boundary between the globus pallidus and the putamen is clearly defined on the mean b1000 DWI given its lower signal intensity resulting from short T2*. Thus, to separate these structures, the initial surface model of the globus pallidus was first deformed to the closest edge on the mean b1000 DWI image (Figure 3B). The striatum was then deformed to the closest edge FA + MD composite map while restricting movement into the initial globus pallidus segmentation (Figure 3C). To prevent the left and right thalamus from intersecting along the medial boundary of these structures, both the left and right thalamus are deformed at the same time to the closest edge on the FA + MD composite map (Figure 3D).

**Figure 3.**
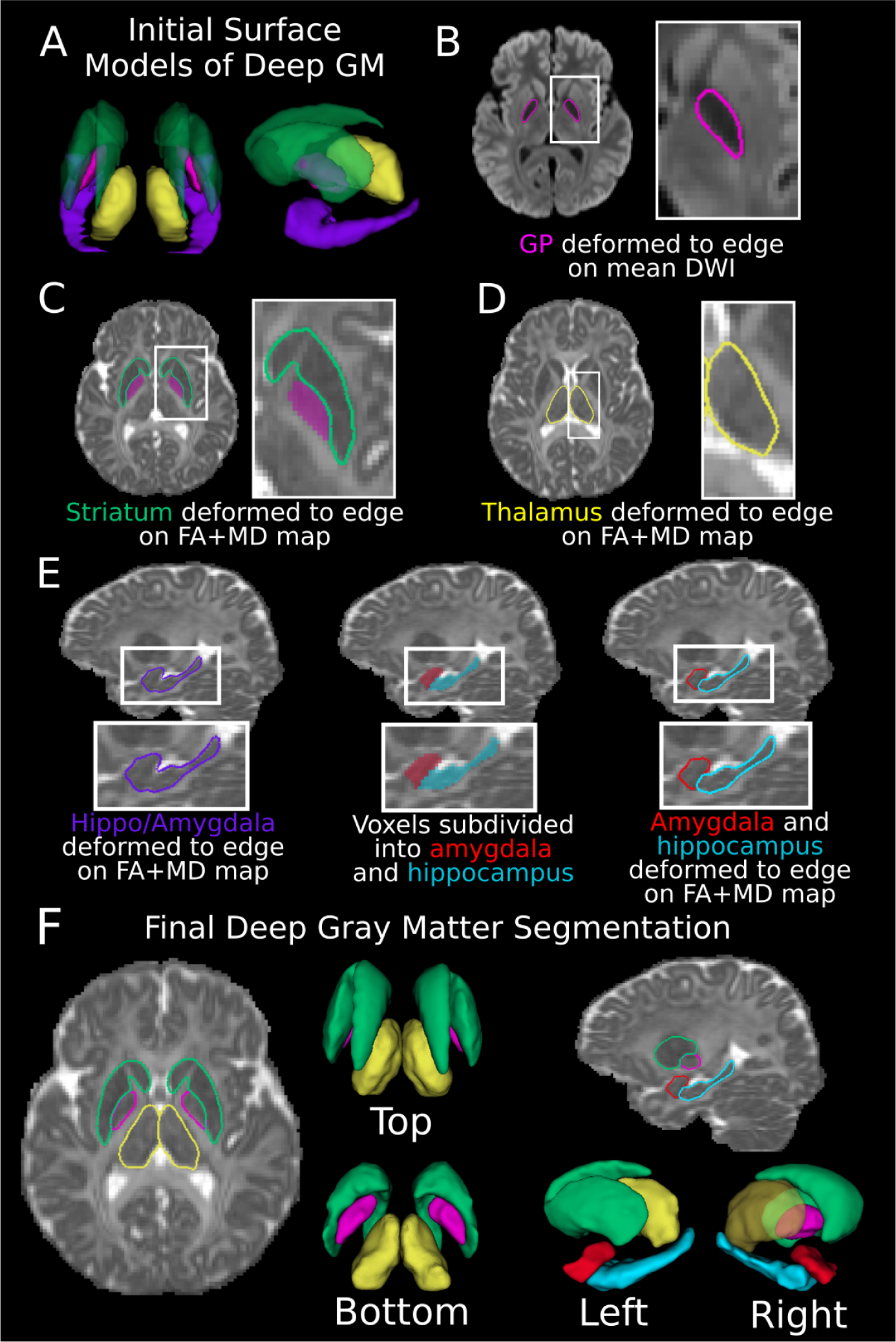
Segmentation workflow of deep GM structure based solely on diffusion tensor imaging visualized on diffusion images acquired with 1.5×1.5×1.5 mm^3^ resolution at 3T. (A) Initial surface models are created from registering a probabilistic atlas to native space. These structures are subsequently deformed one at a time using the mean b1000 diffusion weighted image (DWI), and the FA + MD composite map. (B) First, the globus pallidus (GP, pink) is deformed on the mean b1000 DWI. The remaining structures are deformed subsequently to the nearest edge on FA + MD composite map while restricting surface deformations into the previously segmented structures using the following order of striatum (C, green) then thalamus (D, yellow). Given the lack of contrast between hippocampus and amygdala on DTI, these structures are segmented using multiple steps as follows. (E) First, the hippocampus/amygdala combined surface model is deformed to the nearest edge on the FA + MD map. Voxels within this region are then segmented into amygdala (red), hippocampus (cyan) and cortex voxels based on image intensity of the FA + MD map and registration to a probabilistic atlas (ANTS Atropos). Surface models are then created for each of these voxel-based segmentations and deformed to the nearest edge on the FA + MD composite map. (F) Visualization of final deep GM segmentations for the globus pallidus, striatum, thalamus (top and bottom) and for the left hemisphere for all 5 structures (left and right).

The lateral and medial boundaries of the hippocampus and amygdala are well defined on the FA + MD composite map; however, the boundary between the two structures are not easily discernible. Thus, the hippocampus and amygdala are segmented using multiple steps. First, the hippocampus / amygdala combined mesh is deformed to the closest edge on the FA + MD composite map (Figure 3E). Then a voxel wise label of this region is extracted and then these voxels are relabeled into hippocampus, amygdala and cortex subregions with expectation maximization segmentation (ANTs Atropos (Avants et al., 2011)) using the original hippocampus, amygdala, and cortex probability maps which were registered to native diffusion imaging space. Surface meshes are then extracted for the hippocampus and amygdala regions. Then the hippocampus mesh is deformed again to the closest edge on the FA + MD composite map while restricting the surface deformation into the amygdala label and the amygdala mesh is deformed to the closest edge on the FA + MD composite map while restricting the deformation into the hippocampus and cortex labels. These steps output 3D models of the hippocampus and amygdala where the lateral and inferior/superior boundaries of these structures are primarily determined by the contrast of the FA + MD composite map and the boundary between the structures is primarily determined by the registration of the probability maps.

### 2.4 Comparison of DTI-Only Segmentation to Conventional Methods and Reproducibility of Proposed Method

#### T1-weighted segmentation

To compare the segmentations generated from the proposed segmentation method based solely on DTI images and maps to a more conventional approach, deep GM segmentations were generated on the high resolution T1-weighted images (0.7 mm isotropic) using the standard HCP segmentation pipeline based on FreeSurfer (Glasser et al., 2013). For the University of Alberta test/retest and normative database, T1-based subcortical GM segmentations were generated using FreeSurfer v5.3 (Fischl, 2012). Because the proposed diffusion only segmentation method segments the striatum as a single continuous structure, the T1-based segmentations for the caudate, putamen and nucleus accumbens were combined outputting a single striatum label for each subject in both cohorts.

#### Segmentation overlap comparison

For each subject in the HCP test-retest cohort (first scan only), the T1-based segmentations were interpolated (nearest neighbors) to the diffusion data. Note that the preprocessed diffusion data comes previously registered to the T1w images (mean b0 to T1w image, FLIRT BBR registration (Greve & Fischl, 2009)). As a quantitative comparison between segmentation methods, Dice coefficients (overlap between labels) were calculated for each structure label from the proposed DTI segmentations (voxels contained within each 3D model) to the interpolated FreeSurfer T1w segmentations outputting an inter-method Dice coefficient per subject for the globus pallidus, striatum, thalamus, hippocampus and amygdala keeping left and right separate. The diffusion data available from the University of Alberta normative database was not corrected for phase related distortions (no reverse phase encoding b0 images acquired), thus this data was not included in the segmentation overlap comparison because the distortions would result in a suboptimal registration between the T1-weighted and diffusion images.

#### Structural volume comparison

In addition to Dice scores, regional brain volumes were calculated by summing the voxels included in each segmentation per structure. Differences between the two methods were assessed statistically using paired t-tests for each structure (p < 0.05) using the first scans of the HCP and University of Alberta test-retest data.

#### Segmentation reproducibility analysis

To enable a comparison between the scan 1 and scan 2 segmentations generated from the proposed method, a rigid affine transformation was calculated from the scan 2 mean b0 image to the scan 1 mean b0 image for each subject (Jenkinson et al., 2002) in the HCP and University of Alberta test-retest datasets. The transformation was then applied to the segmentations from scan 2 and nearest neighbors interpolated to native imaging space on the diffusion images of scan 1. Inter-scan Dice coefficients were then calculated between the segmentations from scan 1 and scan 2 for both datasets.

### 2.5 Lifespan Deep Gray Matter Diffusion Measurements

The proposed DTI segmentation method was performed on the diffusion data from the two local lifespan imaging cohorts separately. 3D mesh segmentations were overlaid on the mean b1000 DWI and FA + MD composite map for each subject to qualitatively assess accuracy of the subcortical segmentations. For each subject, MD and FA were calculated as the average from voxels enclosed within each of the 3D mesh structures and volumes were calculated by summing the enclosed voxels of each structure. To compare volumetric trajectories to those from conventional T1-based methods, volumes were generated using FreeSurfer v5.3 (Fischl, 2012) on the lifespan cohorts. Hemispheric differences were assessed for each structure (pairwise t-test, p < 0.05), and because no interhemispheric differences were detected for FA or MD, the values were averaged between hemispheres.

Average MD and FA values were then calculated for each structure (left/right averaged) separately. To assess changes in volume and diffusion parameters related to age and sex, regression modeling was conducted for each lifespan cohort separately as follows. FA, MD, volume (DTI segmentation) and volume (FreeSurfer T1-based) trajectories were estimated separately for each structure with linear, quadratic, and cubic curves; significant curves with the lowest AIC values were considered the best fit (Seabold & Perktold, 2010). In a supplementary analysis of the University of Alberta Normative database, average FA, MD and Volume (DTI segmentation) were calculated separately for males and females for each structure and an independent sample’s t-test (p < 0.001 uncorrected for age and multiple comparisons) was used to assess statistical differences associated with sex. Subsequently the same model fitting procedure was applied separately for males and females to investigate age-related trajectories for each measure by sex.

## 3. Results

### 3.1 Assessment of Deep Gray Matter Segmentations on Native DTI

Based on qualitative assessment for the 1.25 mm isotropic HCP test-retest cohort, segmentations bordered the observable tissue contrast on the mean b1000 diffusion weighted image for the left/right globus pallidus and the FA + MD composite map for the left/right striatum and thalamus (example Figure 4A-C) for all 44 subjects. However, some instances of incorrect segmentations were observed for the hippocampus and amygdala regions. On the HCP test-retest data, hippocampus segmentations were deemed accurate on all but two subjects, where one subject had the bilateral hippocampus segmentations expanded laterally into the cortex and another subject had the same error only on the right hippocampus. Amygdala segmentations in both datasets consistently delineated gray matter voxels; however, the anterior boundary of this segmentation often included cortex due to the lack of contrast between amygdala and cortex on the FA + MD composite map.

**Figure 4.**
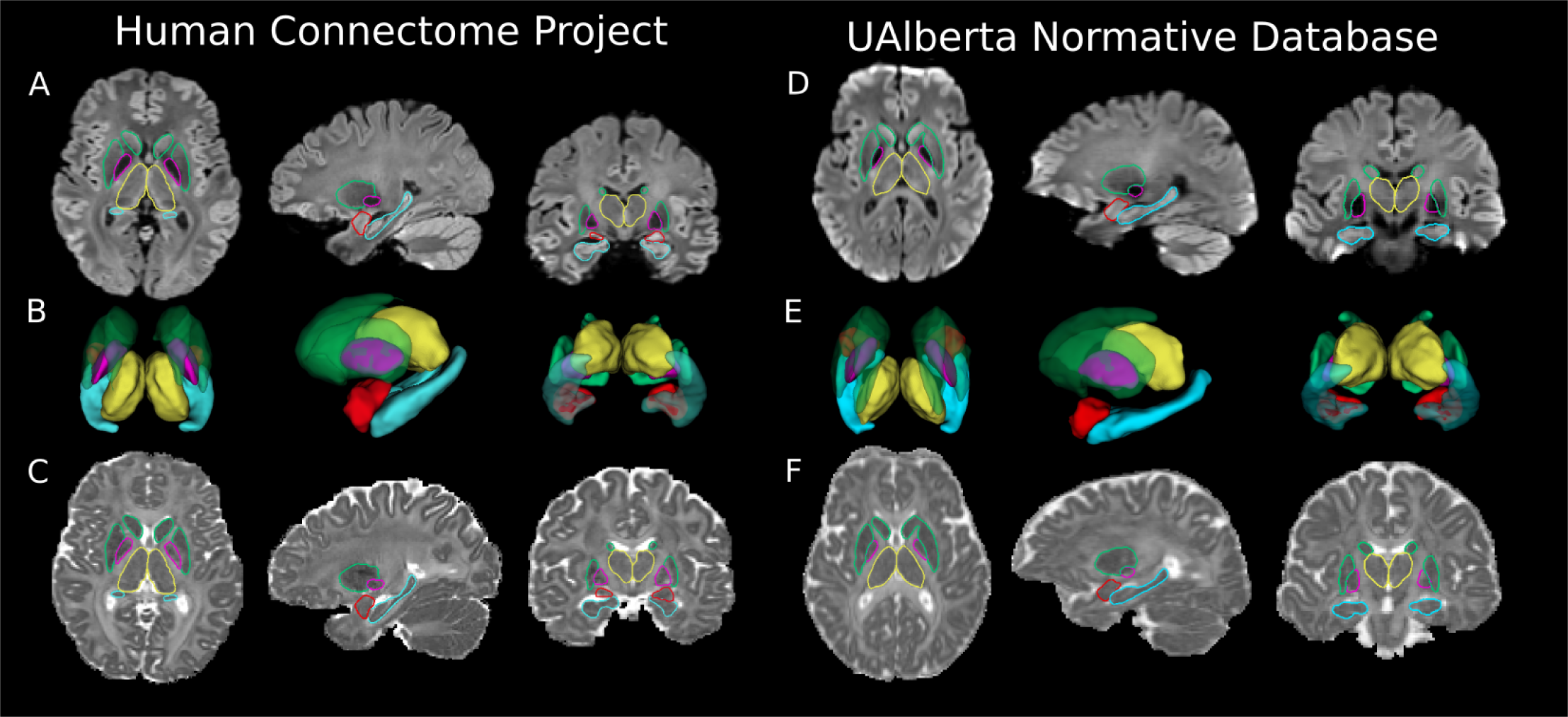
Example segmentation output generated for two participants from the two test/retest datasets used for the validation of the proposed DTI-only segmentation algorithm: A-C (left) - 1.25 mm isotropic diffusion data from the Human Connectome Project (27 year old female) and D-F (right) - 1.5 mm isotropic diffusion data from the University of Alberta normative brain imaging database (33 year old male). Segmentations are displayed for the globus pallidus (pink), striatum (green), thalamus (yellow), hippocampus (cyan) and amygdala (red). Visualizations of slice intersections of deep GM surface models are displayed on (A, D) the mean b1000 diffusion weighted image and (C, F) FA + MD composite map, as well as (B, E) a 3D rendering of the surface models.

On the lower resolution 1.5 mm isotropic University of Alberta test-retest diffusion data, the segmentation algorithm performed equally as well with segmentations of the globus pallidus, striatum and thalamus (example Figure 4D-F) for all 24 subjects bordering the high contrast regions visible in the diffusion images, but with the same errors observed in a subset of the subjects for the hippocampus and amygdala. In this data, the right hippocampus segmentation expanded laterally into the cortex on two subjects with the same segmentation error observed in the bilateral hippocampus segmentations of a separate subject.

### 3.2 Comparison to Standard T1-weighted Segmentation Method

An example of the head to head comparison of the T1-based (0.7 mm isotropic) HCP FreeSurfer segmentation pipeline and the proposed diffusion only method is displayed in Figure 5. Although the majority of the segmentations were in agreement, the standard method produced errors consistently in a few regions that are clearly observed when viewed on the mean b1000 DWI and the FA + MD composite map. The standard method consistently excluded voxels belonging to the globus pallidus in regions adjacent to the posterior limb of the internal capsule (Figure 5A). The standard method also consistently included white matter voxels along the lateral portion of the striatum segmentation adjacent to the putamen and included voxels belonging to the posterior limb of the internal capsule along the lateral portion of the thalamus (Figure 5B). These errors in the conventional method were present in the T1 weighted imaging space where the segmentations were performed (Figure 5C) suggesting that the errors were caused by a lack of GM/WM T1 weighted contrast in these regions and not by the nearest neighbors interpolation to native diffusion imaging space.

**Figure 5.**
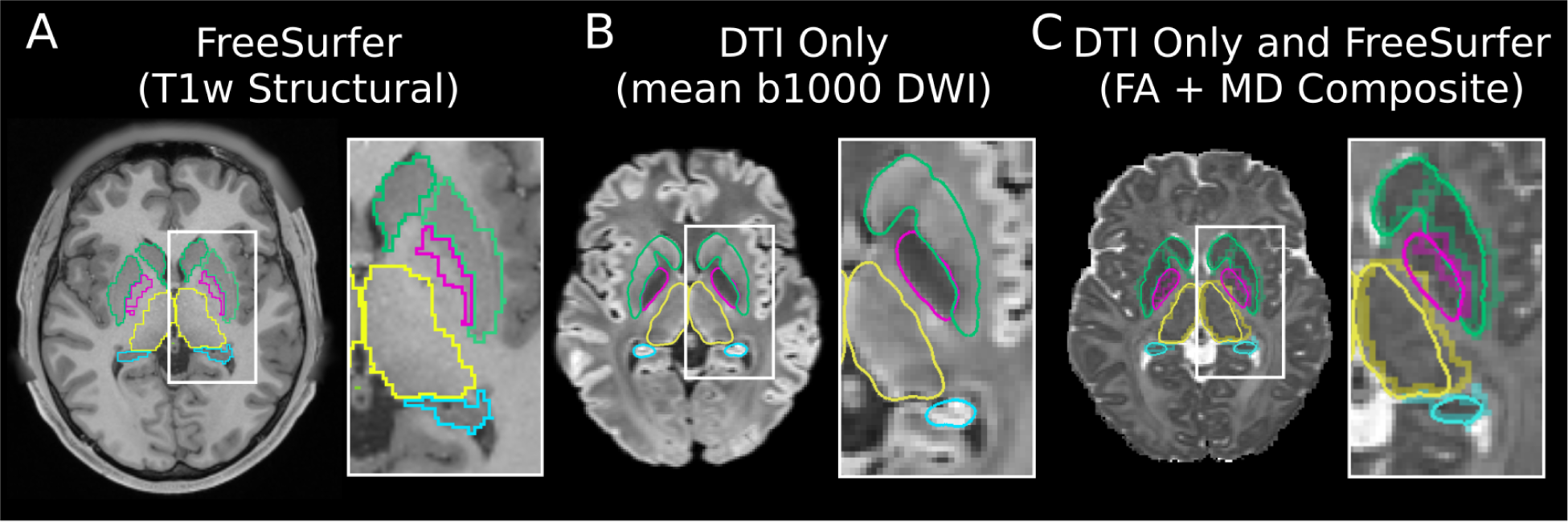
A) Voxelwise segmentation generated from FreeSurfer on the HCP T1-weighted data (22 year old male) overlaid on an axial slice from a T1w structural image. B) Slice intersections of 3D models generated from the proposed DTI only segmentation method overlaid on the mean b1000 DWI from the approximately same slice relative to the T1w image in the same subject. C) Slice intersections from the proposed DTI only segmentation method displayed along with FreeSurfer segmentations (nearest neighbors interpolation from anatomical image to diffusion data) overlaid on the FA + MD composite map. Segmentations are displayed for the globus pallidus (pink), striatum (green), thalamus (yellow) and hippocampus (cyan). In this example, the voxelwise FreeSurfer segmentation poorly delineates the edge of the 4 structures reflecting the poor T1w GM/WM contrast in regions adjacent to these structures.

Group average (HCP test-retest first scan) subcortical GM volumes for the left/right hemispheres for all 5 structures are presented for each method (Table 1) and for the University of Alberta (test-retest first scan) comparison to FreeSurfer T1 segmentation (Table 2). Relative to standard T1-weighted segmentation algorithm, the proposed DTI-only segmentations generated statistically larger volumes (percent difference between 11% and 15%) on the HCP data for 2 of 10 structures (left/right globus pallidus), smaller volumes (percent difference between −28% and −5%) for 7 structures (left/right striatum, left/right thalamus, left/right hippocampus and left amygdala) and no difference in 1 structure (right amygdala). Even with statistically different volumetric measurements, a large proportion of the voxels were in agreement between the two approaches on the HCP data, as evidenced by Dice scores larger than 0.7 in 7 of the 10 structures. The left/right thalamus had Dice scores greater than 0.80 whereas the left globus pallidus and left/right amygdala segmentations had Dice scores lower than 0.7 (Table 1). In the University of Alberta data, the DTI-only segmentations yielded volumes that were statistically lower (percent difference between −30% and −7%) than the T1 Freesurfer segmentations in all 10 structures (left/right striatum, globus pallidus, thalamus, hippocampus and amygdala), likely reflecting the comparatively lower spatial resolution compared to T1w imaging.

**Table 1.** Group average volume measurements for the HCP test-retest cohort extracted per structure using standard 3D T1-weighted automated segmentation (Freesurfer) and the proposed diffusion only segmentation method, along with Dice scores reflecting inter-method voxel overlap.

| HCP Test-Retest Data (n=44, first scan only) |  |  |  |  |  |
| --- | --- | --- | --- | --- | --- |
|  | 3D<br>T1-weighted<br>FreeSurfer<br>(Volume<br>mm<sup>3</sup>) | Proposed<br>Diffusion Only<br>Method<br>(Volume<br>mm<sup>3</sup>) | T1w vs.<br>DTI-Only<br>Percent<br>Volume<br>Difference | Volume<br>Difference<br>(p value) | Inter-Method<br>Dice |
| Left Striatum | 9842 ± 1128 | 8027 ± 869 | -18% | <0.001 | 0.79±0.02 |
| Right Striatum | 10012 ± 1082 | 8030 ± 903 | -20% | <0.001 | 0.80±0.02 |
| Left Globus Pallidus | 1368 ± 253 | 1570 ± 190 | 15% | <0.001 | 0.54±0.26 |
| Right Globus Pallidus | 1427 ± 187 | 1582 ± 184 | 11% | <0.001 | 0.74±0.10 |
| Left Thalamus | 8324 ± 847 | 7011 ± 670 | -16% | <0.001 | 0.82±0.04 |
| Right Thalamus | 7481 ± 746 | 7100 ± 741 | -5% | <0.001 | 0.87±0.02 |
| Left Hippocampus | 4335 ± 437 | 3119 ± 396 | -28% | <0.001 | 0.74±0.02 |
| Right Hippocampus | 4341 ± 475 | 3178 ± 404 | -27% | <0.001 | 0.75±0.03 |
| Left Amygdala | 1529 ± 202 | 1413 ± 188 | -8% | <0.001 | 0.67±0.05 |
| Right Amygdala | 1613 ± 202 | 1641 ± 194 | 2% | 0.116 | 0.69±0.04 |

**Table 2.** Group average volume measurements for the University of Alberta normative database test-retest cohort extracted per structure using standard 3D T1-weighted automated segmentation (FreeSurfer) and the proposed diffusion only segmentation method.

| University of Alberta Normative Database Test-Retest Data (n=24, first scan only) |  |  |  |  |
| --- | --- | --- | --- | --- |
|  | 3D<br>T1-weighted<br>FreeSurfer<br>(Volume<br>mm<sup>3</sup>) | Proposed<br>Diffusion<br>Only Method<br>(Volume<br>mm<sup>3</sup>) | T1w vs.<br>DTI-Only<br>Percent<br>Volume<br>Difference | Volume<br>Difference<br>(p value) |
| Left Striatum | 9694 ± 722 | 7751±540 | -20% | <0.001 |
| Right Striatum | 9196 ± 641 | 7904±487 | -14% | <0.001 |
| Left Globus Pallidus | 1447 ± 150 | 1347±88 | -7% | <0.001 |
| Right Globus Pallidus | 1485 ± 147 | 1372±119 | -8% | <0.001 |
| Left Thalamus | 9010 ± 967 | 6614±509 | -27% | <0.001 |
| Right Thalamus | 8041 ± 852 | 6954±631 | -14% | <0.001 |
| Left Hippocampus | 4752 ± 342 | 3323±270 | -30% | <0.001 |
| Right Hippocampus | 4677 ± 376 | 3256±370 | -30% | <0.001 |
| Left Amygdala | 1660 ± 203 | 1281±237 | -23% | <0.001 |
| Right Amygdala | 1613 ± 162 | 1499±188 | -7% | <0.001 |

### 3.3 Reproducibility Analysis

Using the two test-retest cohorts, volumes and test-retest Dice scores were calculated for all five subcortical structures keeping left and right hemispheres separate (Table 3). The proposed DTI-only segmentation method was highly reproducible in the HCP test-retest diffusion data and the University of Alberta normative data indicated by high Dice scores ranging from 0.88 (left amygdala) to 0.95 (left/right thalamus) in the HCP test-retest data and ranging from 0.79 (left amygdala) to 0.94 (left/right thalamus) in the University of Alberta test-retest data. Paired t-tests revealed no statistical difference (all p > 0.05 uncorrected) in volume measurements between scan 1 and scan 2 for all structures in both the HCP test-retest and the University of Alberta test-retest datasets. Notably, in the retest data, the same segmentation errors noted for the hippocampus and amygdala on the scan 1 data were reproduced in the exact same regions and subjects in both the HCP and University of Alberta data.

**Table 3.** Reproducibility of proposed diffusion only segmentation method over two scans. Group average volumetric measurements for the two separate test-retest cohorts extracted per structure using the proposed diffusion only segmentation method along with inter-scan dice coefficients. No significant pairwise differences (p < 0.05) were observed between scan 1 and scan 2 in either test-retest cohort.

|  | HCP test-retest (n=44) |  |  | University of Alberta test-retest (n=24) |  |  |
| --- | --- | --- | --- | --- | --- | --- |
|  | Scan 1<br>Volume<br>[mm<sup>3</sup>] | Scan 2<br>Volume<br>[mm<sup>3</sup>] | Inter-Scan<br>Dice | Scan 1<br>Volume<br>[mm<sup>3</sup>] | Scan 2<br>Volume<br>[mm<sup>3</sup>] | Inter-Scan<br>Dice |
| Left Striatum | 8027±869 | 7996±887 | 0.93±0.03 | 7751±540 | 7691±578 | 0.91±0.02 |
| Right Striatum | 8030±903 | 8047±916 | 0.92±0.03 | 7904±487 | 7869±578 | 0.91±0.02 |
| Left Globus Pallidus | 1570±190 | 1559±180 | 0.90±0.04 | 1347±88 | 1344±112 | 0.89±0.03 |
| Right Globus Pallidus | 1582±184 | 1588±184 | 0.90±0.04 | 1372±119 | 1378±108 | 0.89±0.04 |
| Left Thalamus | 7011±670 | 6967±651 | 0.95±0.02 | 6614±509 | 6570±597 | 0.94±0.02 |
| Right Thalamus | 7100±741 | 7064±717 | 0.95±0.02 | 6954±631 | 6916±622 | 0.94±0.02 |
| Left Hippocampus | 3119±396 | 3145±396 | 0.90±0.03 | 3323±270 | 3283±361 | 0.87±0.03 |
| Right Hippocampus | 3178±404 | 3159±379 | 0.91±0.03 | 3256±370 | 3259±423 | 0.88±0.03 |
| Left Amygdala | 1413±188 | 1410±180 | 0.88±0.04 | 1281±237 | 1275±212 | 0.79±0.07 |
| Right Amygdala | 1641±194 | 1648±198 | 0.89±0.03 | 1499±188 | 1441±180 | 0.84±0.05 |

### 3.4 Deep Gray Matter DTI Metrics and Volume Versus Age

The diffusion-only segmentations worked well across the lifespan as shown for 3 subjects with very different ages (Figure 6). Notably, in regions with lost signal due to T2* decay observable in older subjects on the mean b1000 DWI, segmentations of the globus pallidus and striatum were also reconstructed accurately (example Figure 6, 82 year old subject). Amongst all subjects in the two lifespan data sets, 5 were excluded from the cohort #1 because of erroneous amygdala segmentations caused by the poor signal around the air tissue interface adjacent to the inferior frontal region of the brain, whereas 1 subject was excluded from cohort #2 due to poor amygdala and hippocampus segmentations. FreeSurfer failed to reconstruct segmentations on T1 images for 3 subjects from cohort #1 which were excluded from further analysis. After excluding subjects, N=357 (37 ± 22 (5 - 90) years; 203 females) remained from cohort #1 and N=164 (30 ± 19 (5 - 74) years; 73 females) remained from cohort #2.

**Figure 6.**
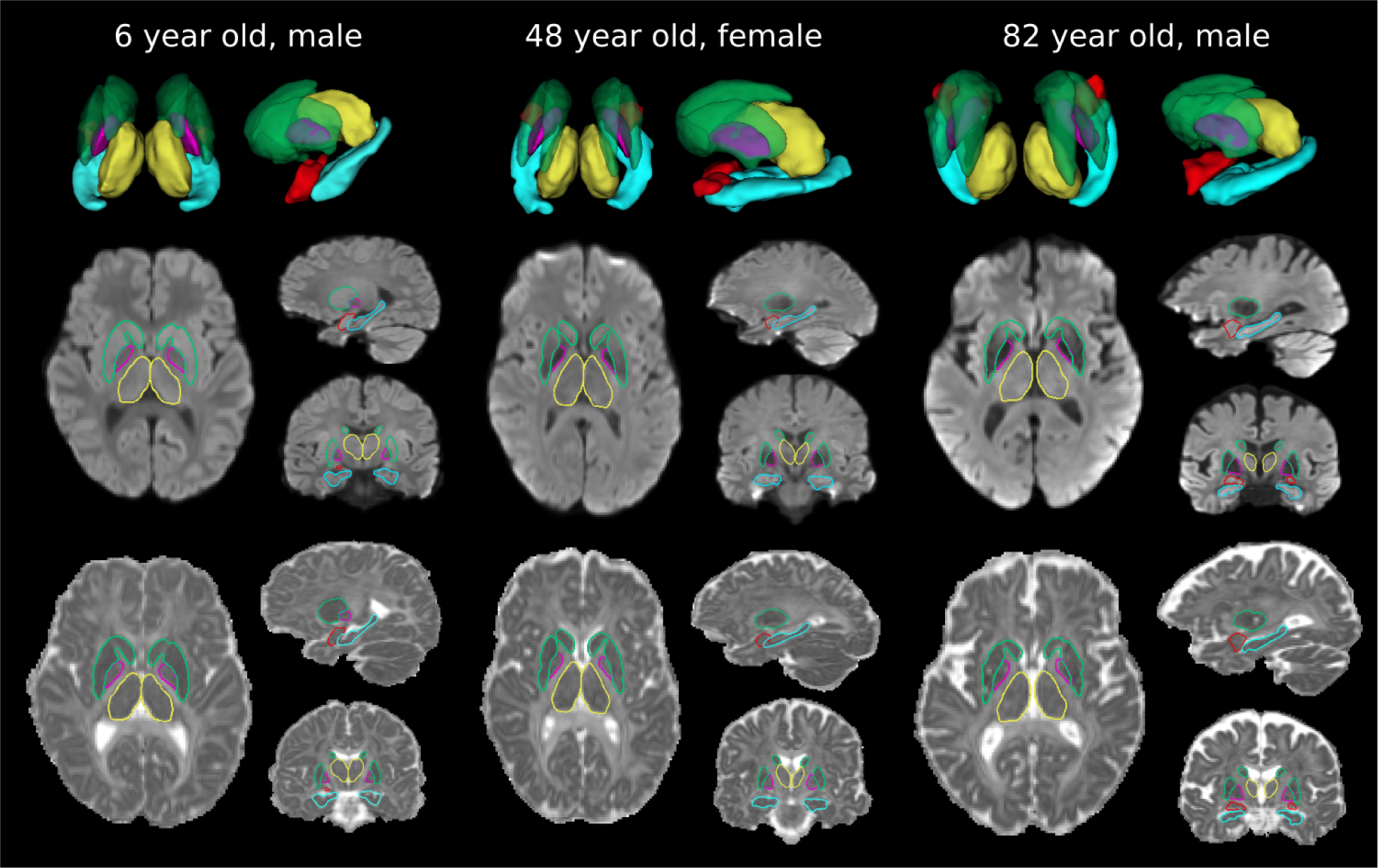
Segmentations generated from the proposed DTI only method displayed for three participants spanning a large age range in the lifespan cohort #1. Segmentations are displayed in 3D (row 1) as well as on slice overlays for the mean b1000 DWI image (row 2) and the FA + MD composite map (row 3). All 5 regions output from the proposed method are displayed including the globus pallidus (pink), striatum (green), thalamus (yellow), hippocampus (cyan) and amygdala (red). Even with substantial subject variability in brain shape, the segmentation algorithm generates structures that align well with the image edges visible on the diffusion images.

Across all subjects from cohort #1, distinct FA values were observed for each subcortical GM structure with higher FA values being observed for the thalamus (0.25 ± 0.01) and globus pallidus (0.23 ± 0.02) compared to the amygdala (0.18 ± 0.02), striatum (0.16 ± 0.01) and hippocampus (0.14 ± 0.01). MD values were lower for the globus pallidus (0.78 ± 0.04 x 10^-3^ mm^2^/s) and striatum (0.81 ± 0.02 x 10^-3^ mm^2^/s) as compared to the amygdala (0.86 ± 0.02 x 10^-3^ mm^2^/s), thalamus (0.87 ± 0.03 x 10^-3^ mm^2^/s), and hippocampus (0.93 ± 0.03 x 10^-3^ mm^2^/s). Cohort #2 (with a slightly different DTI protocol) showed higher average FA values for each structure (thalamus 0.29 ± 0.01, globus pallidus 0.28 ± 0.02, amygdala 0.20 ± 0.01, striatum 0.18 ± 0.01, hippocampus 0.16 ± 0.01) and lower MD values (globus pallidus 0.76 ± 0.03 x 10^-3^ mm^2^/s, striatum 0.77 ± 0.01 x 10^-3^ mm^2^/s, thalamus 0.84 ± 0.02 x 10^-3^ mm^2^/s, amygdala 0.86 ± 0.03 x 10^-3^ mm^2^/s, hippocampus 0.90 ± 0.02 x 10^-3^ mm^2^/s); however, the ordinal relationship between structures remained the same for both cohorts except for the thalamus which had a slightly lower MD compared to the amygdala in cohort #2.

All five deep GM regions showed significantly different FA or MD versus age trajectories (8 non-linear, 2 linear) in the lifespan cohort #1 over 5 to 90 years (Figure 7). The FA age trajectories differed per region with globus pallidus and hippocampus yielding quadratic fits that were flat from ages 5 to ∼60 years of age and then became smaller, amygdala with negative linear correlation, striatum with positive linear correlation, and thalamus with little change (relatively stable cubic trajectory across age). MD values for all structures showed quadratic or cubic U-shaped fits with minima at ∼ 30-40 years where notably the drop of MD during development to the minima was not as much as the greater MD with aging into the elderly, particularly for the thalamus and hippocampus. Cohort #2 with a slightly different DTI protocol, lower upper age, and smaller sample size showed the same general trajectories for FA of the striatum and hippocampus as well as for MD of all 5 deep GM regions, albeit with offset values between cohorts detailed above. The primary cohort #2 differences were in the developmental trajectories for FA of the globus pallidus and the thalamus that increased from 5 to ∼20 years and then leveled off (as opposed to little change) and FA of the amygdala that remained flat from 5-74 years (contrary to a negative linear correlation).

**Figure 7.**
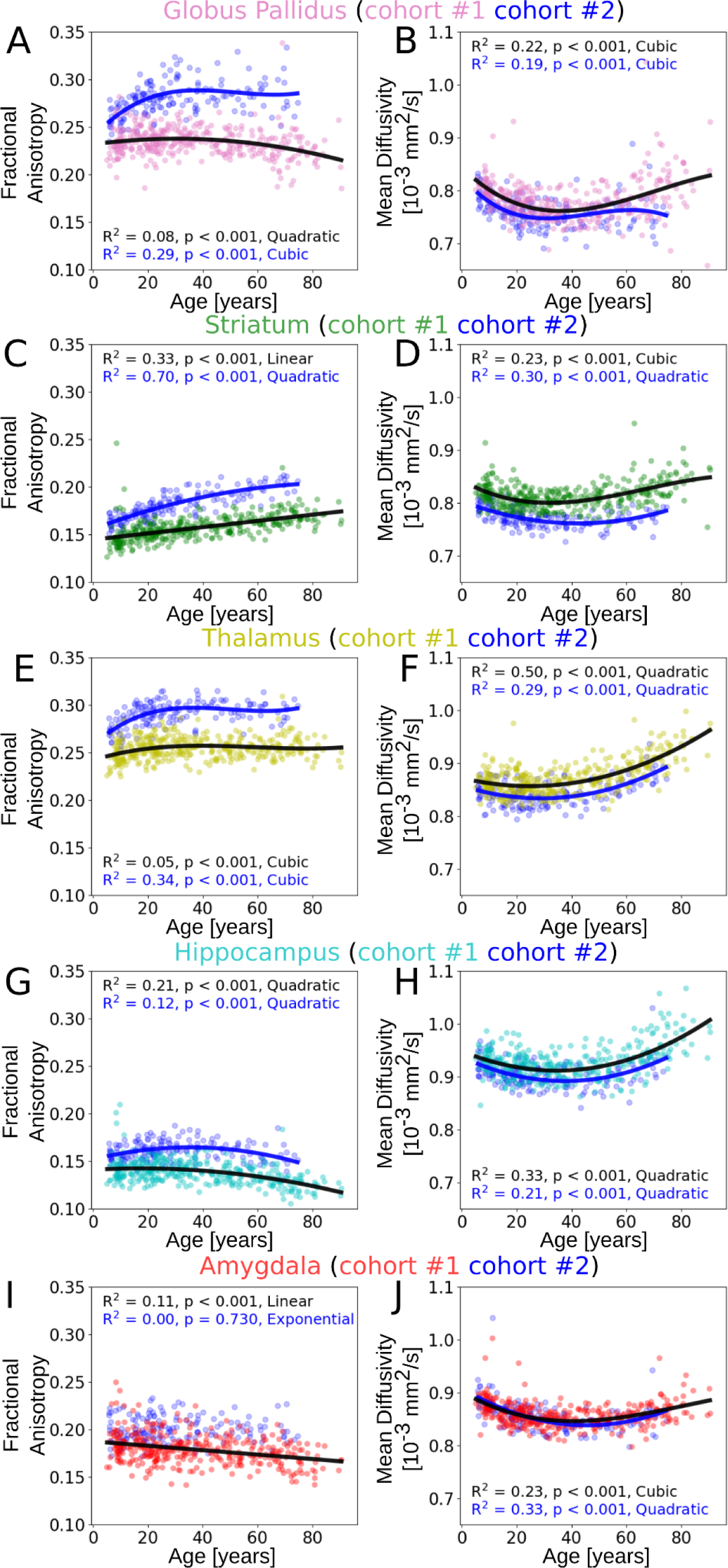
FA (column 1) and MD (column 2) values (left/right averaged) for all subjects from the lifespan cohort #1 (N=365; 5-90 years) plotted by age for the globus pallidus (pink), striatum (green), thalamus (yellow), hippocampus (cyan) and amygdala (red). Scatter plots in those colors are displayed with the best fitting model (black - chosen from linear, exponential, quadratic and cubic fits). In addition, the lifespan cohort #2 (N=164; 5-74 years, with a different 1.5 mm isotropic diffusion acquisition) is plotted for individual DTI values and the best fit model (dark blue points and curve for each structure). FA values remained flat until age 5 to ∼25 years old for the hippocampus and increased linearly for the striatum but showed disagreement between cohorts in the FA trajectories for the globus pallidus, thalamus and amygdala. MD values decreased from childhood to early adulthood (∼30-35 years) and increased thereafter resulting in higher MD values in the elderly compared to childhood.

Different developmental and aging volume trajectories were observed for each structure with marked differences depending on T1 versus DTI segmentation (Figure 8). In the University of Alberta Normative dataset, while using the T1-based FreeSurfer segmentation (Figure 8, column 1), a decreasing non-linear cubic trajectory was observed for the globus pallidus that decreased from 5 to ∼25 years of age and then leveled out thereafter and a decreasing quadratic trajectory was observed for the striatum. The thalamus and hippocampus had trajectories that were flat from 5 to ∼35 years and then declined thereafter, whereas little change was observed through 5 - 90 years for the amygdala. Similar volumetric trajectories were observed when using the DTI only segmentation method for the striatum (linearly decreasing), thalamus (cubic flat from 5 to ∼35 years then decreases) and amygdala (flat quadratic trajectory), whereas the volumetric developmental and aging trajectory of the hippocampus increased from 5 to ∼40 years and then decreased contrary to the T1 approach (flat from 5 to ∼35 years). The biggest difference between developmental and aging trajectories was in the childhood/adolescent age range of the volumetric trajectory for the globus pallidus. With the DTI only method, a cubic trajectory was observed that increased from 5-30 years and then leveled off and decreased thereafter, contrary to the T1 approach (decreased from 5 to ∼25 years then leveled off).

**Figure 8.**
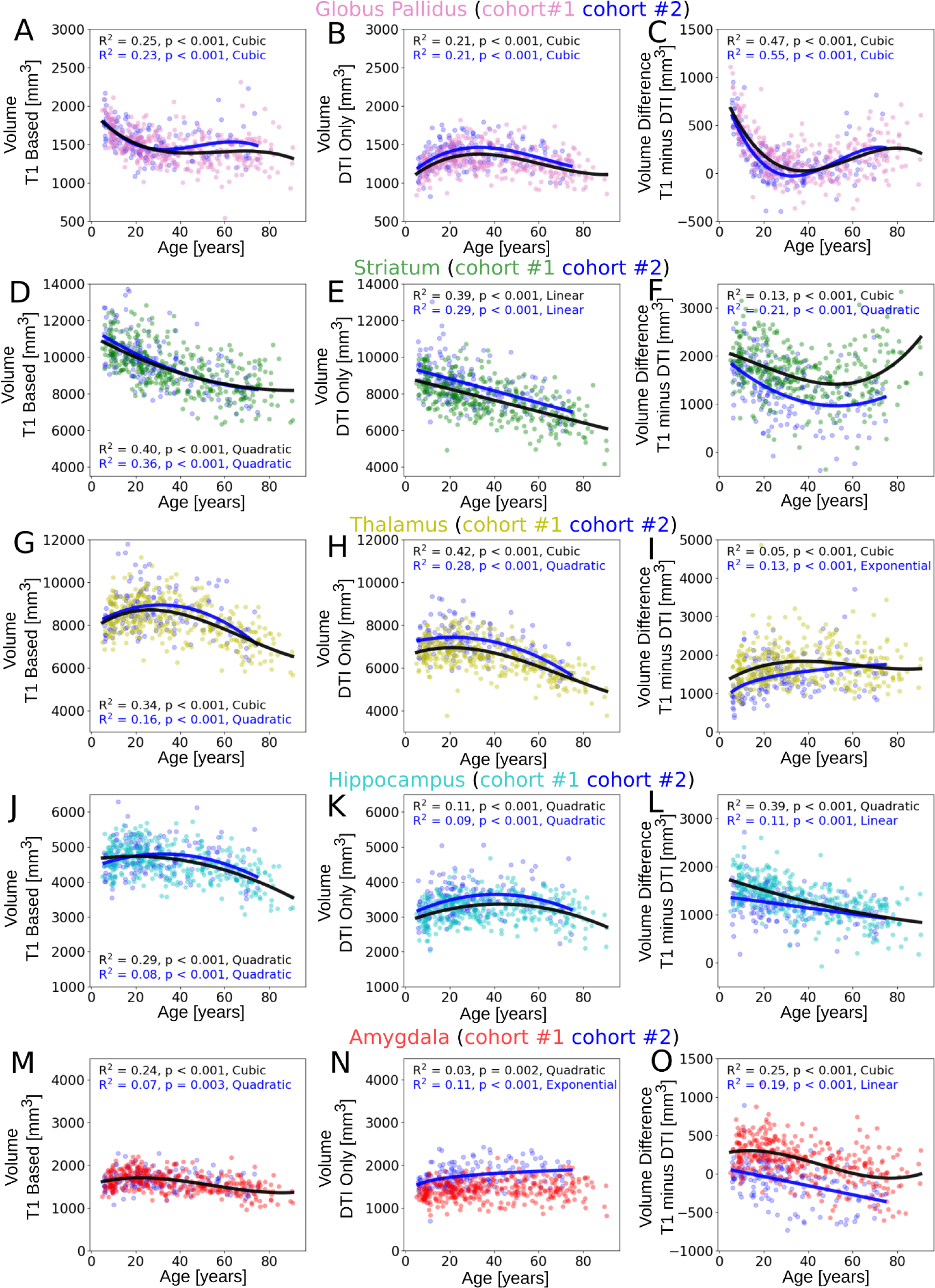
Volume (left/right averaged) calculated on high resolution T1-weighted image (same acquisition between cohorts) using FreeSurfer (column 1) and volume calculated using the proposed DTI only (different 1.5 mm isotropic acquisition between cohorts) segmentation method (column 2) for all subjects from the lifespan cohort #1 (N=365, black curve fits) plotted by age for the globus pallidus (pink), striatum (green), thalamus (yellow), hippocampus (cyan) and amygdala (red) and the lifespan cohort #2 (N=164 all dark blue dots; blue curve fits). The two cohorts show very similar absolute volumes and age trajectories within the same type of image segmentation. However, notable age-related volume differences are apparent between methods (T1-based segmentation volume minus DTI only segmentation volume in column 3). In particular, the globus pallidus T1 segmentation suggests a decrease in volume from 5 to 30 years whereas DTI segmentation suggests an increase over the same age range. The volumes from DTI segmentation were smaller than T1 except the amygdala and 20-50 year age range for globus pallidus.

In both cohorts, average volumes were higher for the T1-based segmentation compared to the DTI only volume across subjects for 4 of the 5 deep GM structures with the following volume differences (T1w volume – DTI volume): globus pallidus (cohort #1: 198 ± 227 mm^3^, cohort #2: 159 ± 236 mm^3^), striatum (cohort #1 1679 ± 572 mm^3^, cohort #2: 1294 ± 581 mm^3^), thalamus (cohort #1: 1711 ± 484 mm^3^, cohort #2: 1442 ± 520 mm^3^), and hippocampus (cohort #1 1323 ± 392 mm^3^, cohort #2: 1200 ± 345 mm^3^). The amygdala had a smaller DTI only volume compared to T1-based segmentation on cohort #1 (71 ± 264 mm^3^) and a slightly larger DTI only volume compared to T1-based segmentation imaging in cohort #2 (−94 ± 251 mm^3^). The T1 segmentation versus DTI segmentation volume differences differed as a function of age, similarly for both lifespan cohorts (Figure 8, column 3). The only two cases with minimal volume differences between methods were the amygdala (Figure 8O) and the globus pallidus but only between the ages of 20-50 years (Figure 8C). At younger ages, the globus pallidus showed a marked reduction in this volumetric T1w-DTI segmentation mismatch from 5 to 20 years (where it became zero) and then the mismatch increased with aging into the elderly. The striatum also showed a U shaped T1w versus DTI volume difference with age. The T1 vs DTI volume difference was smaller with age (although still sizeable) for the hippocampus, but larger with age for the thalamus.

### 3.5 Sex Related Differences in Deep Gray Matter DTI Metrics

Using the proposed DTI only segmentation method, differences associated with sex were assessed for FA, MD and volume using cohort #1 (Figure S1). Developmental and aging trajectories were similar between males and females for all 5 structures for FA, MD and volume. No sex differences were observed for FA (left/right averaged) of the 5 deep GM structures (Figure S1, column 1). MD (left/right averaged) was higher (p < 0.001 uncorrected) in males compared to females for 4 out of the 5 structures (globus pallidus (males: 0.79 ± 0.04 x 10^-3^ mm^2^/s, females: 0.77 ± 0.04 x 10^-3^ mm^2^/s; p < 0.001), striatum (males: 0.82 ± 0.03 x 10^-3^ mm^2^/s, females: 0.81 ± 0.02 x 10^-3^ mm^2^/s; p < 0.001), thalamus (males: 0.88 ± 0.03 x 10^-3^ mm^2^/s, females: 0.87 ± 0.03 x 10^-3^ mm^2^/s; p < 0.001), and hippocampus (males: 0.93 ± 0.03 x 10^-3^ mm^2^/s, females: 0.92 ± 0.03 x 10^-3^ mm^2^/s; p < 0.001), whereas no sex difference was observed for the amygdala. All 5 structures (left/right average) had larger volumes in males compared to females: globus pallidus (males: 1346 ± 181 mm^3^, females: 1230 ± 147 mm^3^; p < 0.001), striatum (males: 8072 ± 1048 mm^3^, females: 7464 ± 1084 mm^3^; p < 0.001), thalamus (males: 6723 ± 798 mm^3^, females: 6360 ± 773 mm^3^; p < 0.001), hippocampus (males: 3369 ± 357 mm^3^, females: 3097 ± 377 mm^3^; p < 0.001) and amygdala (males: 1521 ± 234 mm^3^, females: 1368 ± 200 mm^3^; p < 0.001). Development and aging FA and MD trajectories were similar between males and females for all 5 structures, whereas 4 of the 5 structures had different volumetric developmental and aging trajectories for males compared to females. Specifically, the globus pallidus, thalamus and amygdala appeared to have the same volume for males and females at 5 years of age but increased in males towards 30-50 years of age, and the male striatum reduced in volume at a slower rate relative to females.

## 4. Discussion

### 4.1. Segmentation on DTI Alone

A subcortical gray matter segmentation method is proposed that uses the contrast available on DTI maps and the mean b1000 diffusion weighted image to segment the globus pallidus, striatum, thalamus, hippocampus and amygdala removing the need for the acquisition of or registration to a T1w structural image. Since the segmentation primarily relies on the contrast of the mean DWI and the proposed quantitative FA + MD (x1000) composite map it can be applied on various high resolution DTI acquisitions because FA and MD values for GM, WM and CSF are generally similar between varying protocols. Qualitatively, the proposed segmentation approach had similar segmentation performance when applied to three different DTI acquisitions in test-retest HCP 1.25 mm isotropic data, the University of Alberta Normative database 1.5 mm isotropic test-retest data, and two cross-sectional developing and aging cohorts. The method has been combined with a previously proposed DTI only cortex segmentation method and is available as part of a fully automated diffusion MRI post processing and GM segmentation pipeline (https://github.com/grahamlittlephd/MicroBrain available on publication).

### 4.2. Comparison of DTI Only Segmentation to Standard Approaches

Compared to standard T1-based FreeSurfer segmentation the proposed DTI segmentation method was largely in agreement, indicated by high Dice-scores reported from the HCP test-retest cohort (first scan only) for the striatum (left: 0.79, right: 0.80), thalamus (left: 0.82, right: 0.0.87), hippocampus (left: 0.74, right: 0.75), and right globus pallidus (0.74) segmentations but lower agreement was observed for the left globus pallidus (0.54) and amygdala (left: 0.67, right: 0.69). Two previous studies have proposed DTI only subcortical GM segmentation using machine learning techniques. In a study proposing DORIS, a deep learning based segmentation algorithm, similar FreeSurfer segmentation agreement was reported compared to the current study (globus pallidus: 0.66, striatum (consisting of putamen: 0.78 and the caudate: 0.63, nucleus accumbens excluded), thalamus: 0.78, hippocampus: 0.80 and amygdala: 0.68) (Theaud et al., 2022). The other approach (deepAnat) uses deep learning to create synthetic T1w images from diffusion images and inputs these synthetic images into FreeSurfer. Higher FreeSurfer segmentation overlap was observed using deepAnat (globus pallidus: 0.82, striatum (consisting of the putamen: 0.88, caudate: 0.88 and nucleus accumbens: 0.74), thalamus: 0.91, hippocampus: 0.88 and amygdala 0.82) (Li et al., 2023) compared to the proposed method of the current study. The higher deepAnat Dice-scores compared to the proposed method are not surprising given that deepAnat aims to recreate the same T1w contrast used by FreeSurfer. However, as shown in Figure 5, the T1w contrast may not be ideal for segmenting subcortical GM especially in regions around the internal/external capsules, whereas the DTI maps may provide advantageous contrast for delineating these structures (Figure 1). Because the proposed method uses an interpretable segmentation algorithm, one clear advantage of the proposed method over deep learning based approaches is that it can be intuitively “debugged” when errors arise. Thus, future improvements can be made to the technique without requiring retraining and/or large datasets. Importantly, although all methods here use FreeSurfer output as a “gold standard” segmentation comparison, FreeSurfer itself can generate errors visible when segmentations are overlaid on DTI maps, which is an alternative explanation to imperfect segmentation Dice-scores reported with all previous DTI based segmentation methods. The proposed method was intentionally developed independent of other segmentation approaches, whereas previous deep learning based approaches use FreeSurfer segmentation output for model training and testing (e.g. Billot et al., 2023; Theaud et al., 2022) or use synthetic T1w images as input into FreeSurfer (Li et al., 2023), thus these techniques may be capable of generating the same segmentation errors generated by FreeSurfer.

In the HCP test-retest cohort, lower globus pallidus volumes were measured by T1-based FreeSurfer segmentation for the globus pallidus compared to the proposed method, whereas higher volumes were measured for all other structures. In the test-retest University of Alberta Normative Database higher volumes were measured with FreeSurfer compared to the proposed method for all five subcortical GM structures. The difference in the globus pallidus segmentation was likely a result of the blurred GM/WM T1 contrast adjacent to the internal capsule in the HCP dataset compared to the University of Alberta T1w images where the boundary of the globus pallidus was slightly more visible. The larger T1-based FreeSurfer volumes compare to the proposed method could be explained by many factors, two of which are: 1) imaging resolution of the T1w imaging is far superior to that of the diffusion images causing thinner regions and boundary voxels to be included for a given structure and 2) the poor T1w image contrast causes labels to reach past their true extent, a problem that is mitigated by the contrast available in the diffusion images and maps. Future work will aim to better understand the impact of the diffusion/T1w contrasts on image segmentation with the aim of matching imaging resolution between modalities, a notably challenging problem given the difference in noise levels and effective resolution (amongst many others) between T1w and diffusion imaging techniques.

### 4.3. Deep GM Developmental and Aging DTI Trajectories

Few studies have examined DTI lifespan (excluding the first 5 years) trajectories of the 5 deep GM structures targeted in this study. The proposed DTI only segmentation method enabled this analysis on 529 locally acquired subjects spanning 5-90 years from two different lifespan cohorts. MD and FA values extracted per structure across age were in the expected range for GM (FA: ∼0.15 to ∼0.3, MD: ∼0.75 to ∼1.0×10^-3^ mm^2^/s) and differed across structures suggesting that the diffusion tensor is sensitive to changes in microstructure. Relative to the DTI development and age trajectories of white matter across the lifespan (Lebel et al., 2012; Westlye et al., 2010)(for review see, Lebel et al., 2019), subcortical GM trajectories were more subtle across 5-90 years. In general, MD versus age yielded a U-shaped trajectory which became smaller until mid-adulthood and then increased thereafter for all structures. The MD minima of ∼20 to ∼40 years is comparable to the MD minimum observed for WM tracts across the lifespan (Lebel et al., 2012). In GM, decreases in MD during the first third of life may reflect one of many developmental processes resulting in increased barriers to diffusion (e.g. dendritic arborization, axonal ramification, synaptogenesis, and glial proliferation), whereas the subsequent increases in MD during aging may be associated with the loss of such barriers (e.g. cell loss). FA trajectories showed some increases during development for 4/5 structures (except amygdala) in line with what is seen for white matter tracts, albeit by a lot less, but the subsequent decrease with aging to yield an inverted U trajectory was only observable for globus pallidus and hippocampus. The amygdala showing a steady linear decrease in FA, but given the myriad of cells composing this structure (Yu et al., 2023) a physiological cause of such a trajectory is unclear. In contrast, the FA of the striatum increased with age over the entire lifespan in both cohorts, in agreement with other aging studies reporting linear increases in FA for the putamen (part of the striatum)(Abe et al., 2008; Pal et al., 2011; Wang et al., 2010), but an observation not seen in WM (Lebel et al., 2012). During healthy aging the striatum increases in iron accumulation resulting in faster T2* decay (Treit et al., 2021) and a lower diffusion MRI SNR in this structure. Lower SNR resulting from faster T2* results in the measurement of a higher FA and marginally lower MD (Rulseh et al., 2013), a well known effect when measuring the diffusion tensor in low SNR environments (Jones & Basser, 2004). Thus, the linear developmental aging FA trajectory observed for the striatum may reflect an increase in iron accumulation rather than other underlying microstructural change. To avoid the SNR aging confound in the future, the proposed segmentation method could be leveraged to automatically exclude regions of the striatum with this low signal intensity measured on the mean b1000 DWI.

Between cohorts, the globus pallidus and thalamus had FA trajectories that were similarly flat from ∼20 years onward, but differed substantially between the two lifespan cohorts from 5 to ∼20 years, where both structures showed increasing FA trajectories in cohort #2 but were largely flat in cohort #1. In addition, FA values were consistently higher for all subjects in the cohort #2 compared to the cohort #1. The differences observed between cohorts for FA and MD are most likely not a result of differing segmentations because similar development and aging volume trajectories were observed for cohort #1 and #2 for all five structures (see below). Notably, the diffusion acquisition was different between the two cohorts with the number of diffusion weighted images, zero filling interpolation (which may impact image denoising) unmatched between protocols. In a separate experiment containing 6 participants scanned once with each imaging protocol, lower FA and higher MD values were observed across all 5 structures for the cohort #1 protocol (data not shown), replicating the gross differences observed between the cohorts. This may reflect a larger noise level in the cohort #2 data resulting from poor performance of the denoising algorithm on the on-scanner zero-filled interpolated in-plane imaging data. This noise level difference may also explain the FA increase observed between 5 and ∼20 years in cohort #2, but not cohort #1 for the globus pallidus and thalamus, structures that have similar R2* trajectories over this time period (Treit et al., 2021). Although the majority of the DTI developmental and aging trajectories were similar between cohorts across the majority of the age range, the differences between 5 to 20 years for the globus pallidus and thalamus demonstrate that diffusion imaging acquisition parameters (even on the same scanner) and denoising performance can have a large impact on the measured tensor parameters and alter one’s interpretation if not taken into consideration.

### 4.4. Developmental Alterations in Deep GM DTI

Developmental changes measured by DTI for multiple deep GM structures have been reported previously in studies using manually defined ROIs but excluded the hippocampus and amygdala (Lebel et al., 2008; Pal et al., 2011; Snook et al., 2005). In general, increasing FA and decreasing MD was observed over childhood and adolescence for varying basal ganglia structures (putamen, caudate, globus pallidus, and thalamus) (Lebel et al., 2008; Pal et al., 2011; Snook et al., 2005) with rates of change similar to trajectories from the current study for cohort #2 (globus pallidus and thalamus) and cohort #1/#2 (striatum). The human striatum primarily consists of medium spiny projection neurons and their dendritic spines (Zhou, 2020), thus in development, in addition to iron accumulation, an increase in FA and decrease in MD may reflect an increasingly organized development of dendritic spines throughout adolescence. The increases in FA observed for the globus pallidus observed in previous studies (Lebel et al., 2008; Pal et al., 2011; Snook et al., 2005) and cohort #2 in the current study may reflect the increasing iron accumulation in this structure (as previously discussed). Given the globus pallidus is composed of large pallidal neurons accompanied by large dendritic arborizations and is traversed by myelinated axons of the strato-palidonigral bundle (Yelnik et al., 1984), decreases in MD during development could reflect either increasing myelination and/or alterations to the dendritic and cellular organization during this time period. Interestingly, in cohort #1, no increase in FA but a decrease in MD was observed for the globus pallidus over development, suggesting that the changing DTI measures over development reflect an increase to the hindrance of diffusion rather than an increasingly organized microstructure. The thalamus is composed of multiple nuclei consisting primarily of cell bodies with a sparse number of myelinated fibers that separate nuclei and projection fibers that penetrate its lateral boundaries, hence the source of increasing FA during development observed in cohort #2 and in previous studies may reflect the myelination of these fibers in development. Alternatively, no developmental change in MD or FA was observed for the thalamus in cohort #1 suggesting little to no microstructural maturation of this structure in development. The amygdala and hippocampus were not measured in earlier 1.5T DTI deep GM development studies with limited spatial resolution (3 mm thick slices)(Lebel et al., 2008; Pal et al., 2011; Snook et al., 2005) but their measurement was enabled by the proposed segmentation method and the higher resolution diffusion acquisition used in the current study. Hippocampus MD got smaller over development in both cohorts of the current study, whereas FA was unchanged over development (cohort #1) and got marginally larger (cohort #2). The FA and MD developmental trajectories were similar to those measured from a high-resolution 1 mm isotropic diffusion acquisition specifically designed for the hippocampus and acquired on the same cohort #2 subjects from the current study (Solar et al., 2021). High-resolution developmental MD and FA trajectories more closely matched those measured from the cohort #2 acquisition in the current study supporting the notion of increased FA in development which may relate to increased axonal packing and/or myelination (Coras et al., 2014; Shepherd et al., 2007). Diffusivity as measured by the apparent diffusion coefficient (ADC) has been observed to decrease for the amygdala plateauing at ∼10 years (MacEachern et al., 2020) earlier then the MD plateau observed in the current study (cohort #1 and cohort #2) into young adulthood. A shallow linear decreasing trajectory was observed across development for FA of the amygdala in cohort #1 but not cohort #2, suggesting that the DTI changes observed over this time period are not related to changes along a single axis of the diffusion tensor, whereas previous work has suggested that diffusion along the primary diffusion tensor direction is associated to fibers passing through the amygdala (Solano-Castiella et al., 2010). Taken together, these studies suggest that DTI is sensitive to continued development of these structures throughout childhood and into late adolescence towards young adulthood, contrary to previous work suggesting deep GM FA and MD trajectories level off prior to 4 years old (Shi et al., 2019).

### 4.5. Aging Related Changes in Deep GM DTI

In middle adulthood, MD trajectories modeled for both cohorts from the current study for all five structures level off and begin to increase into later adulthood. The inversion of deep GM MD trajectories were not reported in previous studies that investigated younger cohorts (max age 40 years (Faria et al., 2010) and 52 years (Pal et al., 2011)), an advantage of the current studies’ larger age span. Mid life FA trajectories either declined (globus pallidus (cohort #1), hippocampus (cohort #1 and #2), amygdala (cohort #1)) or plateaued (globus pallidus (cohort #2), thalamus (cohort #1 and #2) in all structures but the striatum which showed a steady linear increase (cohort #1 and #2) throughout middle adulthood. Increases in MD trajectories were observed after ∼30-35 years in all 5 deep GM structures with the most prevalent increases observed in the thalamus and hippocampus. Importantly, the entire medial boundaries of these two structures are adjacent to CSF, thus age related atrophy of these structures may increase the proportion of partial volume measurement of CSF, hence increasing MD. In a high-resolution (1 mm isotropic) DTI lifespan study of the hippocampus (same cohort #2), no significant increase in MD was observed in adulthood, however the segmentation protocol did not include the subiculum a region bordering CSF over the anterior-posterior extent of the hippocampus, suggesting that the increase in MD in the current studies’ hippocampus trajectories were at least partially driven by regions prone to CSF contamination. Interestingly, even the globus pallidus, a region encapsulated by surrounding white matter and the putamen, also showed an MD increase in adulthood. Previous findings have observed larger globus pallidus MD values in adult populations compared to young adults using standard DTI acquisitions (Pfefferbaum et al., 2010) and CSF suppressed FLAIR DTI (Bhagat & Beaulieu, 2004) suggesting that the observed increase in MD observed in the current study is not solely due to CSF partial volume measurement.

Decreasing or constant FA trajectories were observed into adulthood of 4 of the 5 deep GM structures with the exception of the striatum which had a steady linear increase in FA throughout life. Aging associated FA increases in the striatum (putamen and/or caudate) have been consistently reported in both lifespan studies (Hasan et al., 2008, 2009; Pal et al., 2011; Wang et al., 2010) and cross-sectional comparisons between old and young cohorts (Bhagat & Beaulieu, 2004; Pfefferbaum et al., 2010). This increase in FA has been linked to increasing iron accumulation in this structure (Rulseh et al., 2013), however FA increases in the striatum into adulthood were also observed in cohort #1 which is speculated to be less affected by R2* compared to cohort #2 for the globus pallidus and the thalamus (as previously discussed). Thus, another explanation of increasing FA with age in the striatum may be the targeted loss of neurons or particular dendrite connections (Hasan et al., 2008, p. 200). The remaining 4 structures either had no change into adulthood (globus pallidus (cohort #2), thalamus (cohort #1 and #2)) or decrease into late adulthood (globus pallidus (cohort #1), hippocampus (cohort #1 and #2), amygdala (cohort #1)).

Decreases in FA were not observed in previous studies that did not contain elderly participants (max age 40 years (Faria et al., 2010) and max age 52 years (Pal et al., 2011)). In the high-resolution hippocampus lifespan study including lifespan cohort #2, decreased FA was also observed, indicating that this age related change is likely not related to partial voluming of the hippocampus with CSF. A variety of factors may explain decreases in FA in these three structures, one such explanation is the demyelination of fibers that traverse all three of these structures (e.g. strato-palidonigral bundle in the globus pallidus, the fimbria/alveus of the hippocampus, and other projection fibers such as the fornix passing through the amygdala (Solano-Castiella et al., 2010)). However, other microstructural alterations (e.g. neuronal loss, decreased axon packing) may also alter tensor parameters as a function of age and thus can not be ruled out.

### 4.6. Development and Aging Volume Trajectories of Deep GM Measured on DTI

Development and aging volume trajectories for 4 (striatum, thalamus, hippocampus and amygdala) of the 5 subcortical GM structures investigated were largely similar when using the proposed DTI only segmentation method on the DTI images compared to the ubiquitous T1-based volumes (FreeSurfer segmentation in this case). However the T1 segmentations yielded consistently larger volumes than the DTI segmentations, as previously discussed. This means that the developmental and aging volumetric interpretation would remain the same regardless of using the proposed segmentation method or FreeSurfer with these data. Notably, much of the lifespan volumetric trajectories agreed between the proposed method and FreeSurfer, for example the large reduction in thalamus volume observed throughout late adulthood and the continued decrease in striatum volume observed across the entire lifespan (5-90 years).

The exception was in the globus pallidus prior to ∼20 years which had a decreasing volume when measured with FreeSurfer on T1w relative to the DTI-only method which had an increasing volume during this age range. It is unclear whether the globus pallidus increases or decreases in volume during childhood/adolescence even on solely T1w studies, with some reporting increases in the volume of the globus pallidus (Dima et al., 2022; Narvacan et al., 2017), while other work suggests steady declines in globus pallidus volume throughout childhood/adolescence (Fjell et al., 2013). The different developmental trajectory observed for the globus pallidus measured on DTI compared to T1-based segmentation in the current study may reflect the improved GM/WM contrast around the internal capsule provided by the FA + MD composite map; however future work is needed to determine factors related to image acquisition that may influence the volumetric measurement of this structure on DTI (e.g. blooming artifact on SS-EPI due to increased susceptibility).

The proposed method also measured markedly similar trajectories on both cohort #1 and cohort #2, demonstrating that the differences in DTI acquisition protocols between the cohorts, that resulted in different FA and MD trajectories, were not an issue for the segmentation algorithm and resulting volumetric measurements. The proposed DTI only segmentation provides an additional avenue for measuring deep GM volume, however more work is needed to fully understand the true volumetric development and aging trajectories of the subcortical GM and the effects differing segmentation algorithms have on their measurement.

### 4.7. Study Limitations and Future Directions

Frequent errors were encountered when segmenting the amygdala using the proposed method. These errors were a result of 1) the anterior boundary separating the amygdala from the cortex was indiscernible on the FA + MD composite map and/or 2) large distortions caused by the adjacent air tissue interface corrupted the image in this entire region. Importantly, even with these errors the volumes of the amygdala were within range of those reported by FreeSurfer, but even still, FA and MD results reported for the amygdala using the proposed method should be interpreted with caution.

The imaging acquisitions used in the current study were notably higher resolutions of 1.5 mm or 1.25 mm isotropic than a typical 2 mm isotropic whole brain diffusion acquisition. In testing on typical 2 mm isotropic diffusion images, the proposed method failed for all structures except the striatum, suggesting that the method has practical limitations that will be reached when applied to lower resolution diffusion data. High resolution diffusion imaging studies in large cohorts are becoming more common, the diffusion protocol for the HC Development (HCP-D) and Aging (HCP-A) cohorts is acquired at a 1.5 mm isotropic resolution, and the proposed methodology will be applicable to these data sets as well (Harms et al., 2018).

Other work proposing whole brain segmentation on native diffusion images have demonstrated the advantage of using these segmentations to improve WM tractography (Li et al., 2023; Theaud et al., 2022), hence the proposed method may also aid in this context. A unique aspect of the proposed method is that it generates 3D meshes of the subcortical GM structures that align with the GM/WM boundaries visible on the diffusion images/maps. Future work will aim at using the generated meshes to incorporate aspects of shape and size of deep GM structures into tractography algorithms, creating more anatomically plausible tractography reconstructions as has been similarly proposed (Yeh et al., 2017).

Modeling the diffusion signal in gray matter is still an open problem given the existence of molecular exchange between cellular and extracellular compartments (Jelescu et al., 2022). In this study, we chose to represent the diffusion signal as a tensor to avoid assumptions about the underlying microstructure of the 5 deep GM structures of interest. In fact, other studies of development (Mah et al., 2017; Zhao et al., 2021) and aging (Guerreri et al., 2019) have proposed using higher order diffusion models to study the deep GM; however to date many of these higher order models are designed for WM and are unsuited to the microstructural environment of GM so developmental/aging trajectories generated by these techniques should be interpreted with caution. Outside of the scope of this work, future studies will explore diffusion modeling techniques that are more tailored to the microstructural environment of the individual deep GM structures.

## 5. Conclusions

A fully automated segmentation method is proposed that is designed to work directly on diffusion images for the purpose of measuring DTI parameters from 5 deep GM structures (globus pallidus, striatum, thalamus, hippocampus and amygdala). The segmentation uses the contrast available on DTI from the mean b1000 DWI and a FA + MD composite map to deform 3D meshes to the boundaries of each of the target structures. Compared to standard T1-based FreeSurfer segmentation the proposed method generated similar segmentations for the 5 deep GM structures and may outperform T1-based segmentation methods due to the excellent GM/WM contrast visible on the FA + MD composite map throughout the internal/external capsule. In two typical development and aging cohorts including 529 participants over the ‘lifespan’ of 5 to 90 years, unique development and aging trajectories were observed for FA and MD of each deep GM structure that were distinct from their observed volumetric trajectories, suggesting that DTI may provide additional information about brain maturation in the deep GM. The automated DTI only segmentation does not require T1 imaging and has clear advantages over T1-based segmentation. Thus, the method will be useful for future studies of the diffusion properties of deep GM in development, aging or clinical populations, where T1 images either add to the total acquisition time or are not acquired routinely.

## Data Code and Availability

The full segmentation pipeline will be made available upon publication on the first author’s github page (https://github.com/grahamlittlephd/MicroBrain). All data from the University of Alberta normative sample (Treit et al., 2023) will be made publicly available following publication embargos. For download links and data access, please email the Principal Investigator, Christian Beaulieu at christian.

## Author Contributions

**Graham Little:** Writing - original draft, Conceptualization, Methodology, Data curation, Formal analysis, Software. **J. Alejandro Acosta-Franco:** Data curation, Analysis **Christian Beaulieu:** Conceptualization, Methodology, Writing - original draft, Writing - review & editing.

## Acknowledgements

This work was funded by grants from the Canadian Institutes of Health Research (CIHR), University Hospital Foundation, and Womens and Childrens Health Research Institute, as well as a salary award from the Canada Research Chairs program (CB) and fellowships from the Natural Sciences and Engineering Research Council of Canada fellowship (GL) and Unifying Neuroscience and Artificial Intelligence in Quebec fellowship (GL).

---

**Figure S1.**
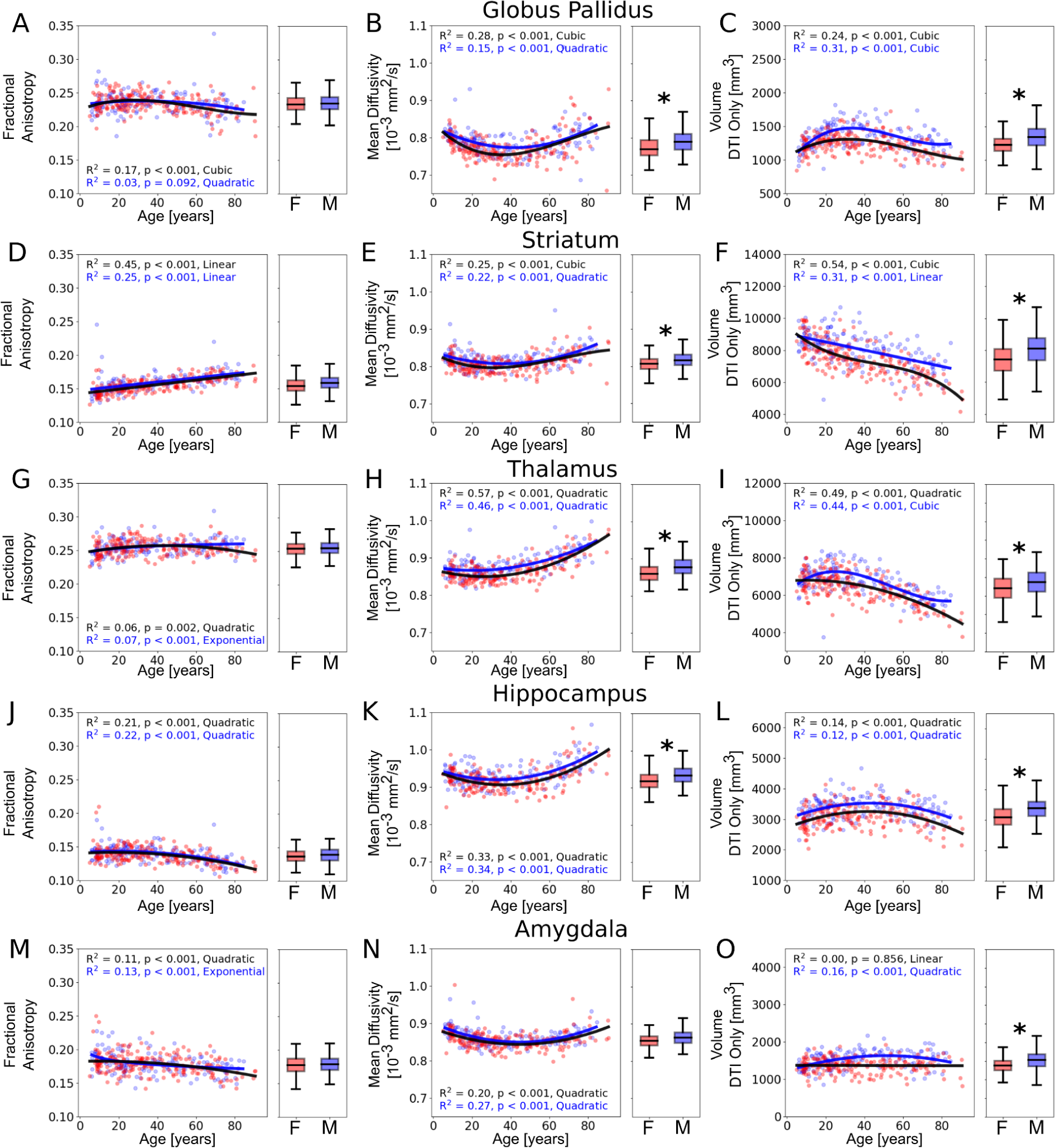
Deep GM FA (column 1), MD (column 2) and Volume (column 3) calculated using the proposed DTI only segmentation method and plotted versus age separately for females (red) and males (blue) for all subjects from the University of Alberta Normative database cohort #1 (N=156 males and 209 females). Trajectories followed the same general trends for males and females for FA, MD and volume. Four (globus pallidus, striatum, thalamus, hippocampus) of the 5 deep GM structures had greater MD for males compared to females (albeit slight) and all 5 structures had a higher volume in males compared to females on average over the shown lifespan (* assessed by t-test p < 0.001).

